# A biophysical dynamic module for the polarisation of auxin efflux carriers PIN-FORMED (PIN)

**DOI:** 10.1101/216275

**Authors:** Valeria Hernández-Hernández, Rafael A. Barrio, Mariana Benítez, Naomi Nakayama, José Roberto Romero-Arias, Carlos Villarreal

**Author notes:** Current Address: Laboratoire de Reproduction et Développement des Plantes, Université de Lyon, ENS de Lyon, UCB Lyon 1, CNRS, INRA.

## Abstract

Intracellular polarisation of auxin efflux carriers is crucial for understanding how auxin gradients form in plants. The polarisation dynamics of auxin efflux carriers PIN-FORMED (PIN) depends on both biomechanical forces as well as chemical, molecular and genetic factors. Biomechanical forces have shown to affect the localisation of PIN transporters to the plasma membrane. We proposed a biophysical module of PIN polarisation that integrates biomechanical, molecular, and cellular processes as well as their non-linear interactions. The module was implemented as a discrete Boolean model and then approximated to a continuous dynamic system, in order to explore the relative contribution of the factors mediating PIN polarisation at the scale of single cell. Our models recovered qualitative behaviours that have been experimentally observed and enabled us to postulate that, in the context of PIN polarisation, the effects of the mechanical forces can predominate over the activity of molecular factors such as the GTPase ROP6 and the CRIB-motif CRIB-motif RIC1.

## INTRODUCTION

Plants are able to generate new organs in an undetermined and plastic manner. Studies on plant development have revealed that, among the many signals that mediate this plasticity, the phytohormone auxin plays a central role regulating basic processes such as cell division, elongation, and differentiation in a concentration-dependent manner (Habets and Offringa, 2014; Zazimalova et al., 2014). Auxin is transported cell-to-cell in a polar manner generating auxin gradients through the tissues that interact with gene regulatory networks (Taiz and Zeiger, 2010) However, it is yet not fully understood how plants dynamically regulate and couple auxin polar transport and perception at the cell, tissue, and organ levels.

At the molecular level, several protein families have been shown to transport auxin into and out of the cells in the model plant *Arabidopsis thaliana* (Feraru and Friml, 2008). Among them, members of the family of efflux carriers PIN-FORMED (PIN) have been well studied. Arabidopsis posses eight different PIN transporters, all of them, except PIN5 and 8, localise at the plasma membrane (Feraru and Friml, 2008). It has been shown that these proteins are actively sorted to particular domains of the plasma membrane – i.e., they are polarised – in a cellular-dependent context and that they can direct auxin effluxes (Blilou et al., 2005; Wisniewska et al., 2006; Feraru and Friml, 2008). *pin* loss-of-function mutations alter the morphology of shoot and root apical meristems (SAM and RAM, respectively) (Okada et al., 1991; Blilou et al., 2005).

The study of PIN polarisation dynamics has revealed some of the molecular and biophysical regulators. Among the molecular factors implicated is the family of GTPase proteins called Rho of Plants (ROPs), which inhibit the formation of clathrin-coated vesicles in which PINs (located at the plasma membrane) are endocytosed (Kitakura et al., 2011; Chen et al., 2012; Lin et al., 2012). This pathway is thought to respond to the auxin signal through the putative auxin membrane receptor AUXIN BINDING PROTEIN 1 (ABP1) (Paciorek et al., 2005; Robert et al., 2010; Chen et al., 2012; Lin et al., 2012, although see Gao et al., 2015; Michalko et al., 2015; Jásik et al., 2016 for contradictory results). Mechanical forces acting at the tissue level, for example deformation (expansion, compression, and shape change), and the tension in the cell wall and the plasma membrane, are able to regulate PIN polarisation patterns (Heisler et al., 2010; Feraru et al., 2011; Nakayama et al., 2012; Braybook and Peaucelle, 2013; Zwiewka et al., 2015). Taken together, our current knowledge suggests that multiple physico-chemical factors act at different spatio-temporal scales to regulate the polarisation patterns of the PIN auxin efflux carriers.

Despite these advances, how the biomechanical forces and molecular genetic factors are orchestrated in PIN polarisation dynamics has not been thoroughly studied. Some models have tried to explain the PIN polarisation based on the interactions among molecular factors alone (Jönsson et al., 2006; Smith et al., 2006; Wabnik et al., 2011), while others have taken into account the biomechanical forces (Heisler et al., 2010). Very few studies have integrated both types of regulators, as well as their feedbacks, to understand how PINs polarise (Newell et al., 2008). Here we used a qualitative network model to integrate the molecular genetic factors with the mechano-sensitive cellular processes to study PIN polarisation at the cellular scale. Our network can be considered as a biophysical dynamic module (*sensu* Newman and Bhat, 2009; Newman et al., 2009) since it incorporates molecular, cellular and mechanical processes, and their feedbacks, that have been reported to affect PIN polarisation. Compared with other studies that take into account the interactions between molecular and mechanical forces during PIN polarisation, the theoretical framework put forward by Newman and collaborators considers that physical factors “impart a discernable and at least semiautonomous role to the functions of the gene products” (Newman and Bhat, 2009). As it is discussed in the following pages, the results of our simulations support a discernable role of the mechanical tension of the plasma membrane in the polarisation dynamics of PIN transporterr from that of the ROP6 GTPase.

Following a protocol established by Azpeitia and coworkers (2014) to study gene regulatory networks, we initially modelled the dynamical module as a Boolean network to explore the qualitative logic of the interacting components. We then approximated the Boolean model to a continuous system to infer the relative role of particular factors. For example, we analysed the impact of *ROP6* loss-of-function mutation on PIN localisation in the context of our biophysical module. While, the Boolean approach provided valuable insights into the regulatory interactions within the system, the continuous approach reliably recovered some of the most relevant features of PIN polarisation dynamics. Our simulations rendered testable predictions, such as the relative roles that the mechanical factors play in establishing PIN polarisation patterns. Furthermore, our models also underscored several empirical gaps in the mechanisms involved in PIN polarisation.

## METHODS AND MODEL SPECIFICATION

### Construction of the biophysical dynamic module

We have incorporated the experimental findings for the factors affecting PIN polarisation up to the literature published in August 2016. The criterion we used to include a factor in our module was that its physical, chemical and/or genetic disruption affected either the localisation of PIN proteins to the plasma membrane, its polarisation patterns, or both. Thus, for example, the D6 PROTEIN KINASE (D6PK) was not taken into account because it acts as regulator of the activity of PINs as auxin carriers, but not of the intracellular localisation of PINs (Habets and Offringa, 2014; Zourelidou et al., 2014). From this literature we constructed an interaction network for a biophysical dynamic module for PIN polarisation (*sensu* Newman and Bhat, 2009). This module accounts for cell autonomous processes for PIN intracellular polarisation and includes biomechanical factors (e.g., the mechanical tension of the plasma membrane), cellular processes (e.g., exocytosis), and molecular interactions (e.g., PIN phosphorylation by the serine/threonine protein kinase PINOID, PID). We described the factors that we selected as well as their interactions with one another in the “Biophysical dynamic module” subsection of the Results.

### Boolean model

We implemented a Boolean network model to study how the different factors included in the biophysical module interact with each other in the context of PIN polarisation dynamics. This is one of the simplest formalism to study the non-linear interactions of agents constituting regulatory networks. It provides meaningful qualitative information about the intricate mechanistic models of a biological system and, therefore, it has been used to describe diverse biological problems (Villarreal et al., 2012; Davidich and Bornholdt, 2013; Fumia and Martins, 2013; LaBar et al., 2013; Davila-Valderrain et al., 2016). In this type of models the state variable of each node is discrete and takes a value of 0 (i.e., inhibited, below a given threshold, OFF or inactive) or 1 (i.e., activated, above a given threshold, ON, present). Nodes can represent genes, proteins, RNA, etc., and links represent positive or negative regulation between pairs of nodes. Each node's state is specified by a function that depends on the state of its regulators at a previous time *t*, as in:

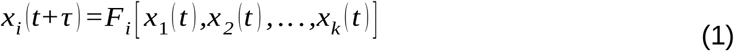

where 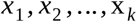 are the regulators of node 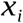 and 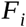 is a discrete function representing a logical proposition formalised in terms of algebraic rules of Boole´s axiomatics. These rules were constructed on the basis of the reviewed experimental evidence for the plant model *A. thaliana* and correspond to the links depicted in Figure 1 and 2. A Boolean network has 2^n^ (n being the number of nodes) possible states of activation/inhibition meaning that the system's dynamics is deterministic. A boolean attractor is a vector composed of stationary values of activation and inhibition of the dynamical mapping that is determined by the condition 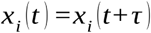 and that defines a homeostatic state of the system. An exhaustive exploration of the attractors of the system was performed by a synchronous evaluation of the logical rules of each node. For that purpose, we employed the BoolNet package of R (version 3.2.4) (the codes for this and the continuous models are freely available in the GitHub link: https://github.com/hhdez/Biophysical-dynamic-module-for-PIN-polarisation). Two methods were used to test the validity of the Boolean model. The first one was the simulation of knock-out (KO) and overexpression (OE) mutants, performed by fixing the activation state of the altered node 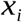 to 0 or 1, respectively. In these simulations we and analysed whether the activation state profiles of the altered attractors corresponded to activity profiles experimentally observed for each mutant. Because of the discrete nature of our model, the comparisons between the attractors and the experimentally reported results are qualitative (see a previous similar analysis in Espinosa-Soto et al., 2004). The second way consisted in the application of the standard robustness test “shuffle” for stochastic perturbation available in the BoolNet package. This method consists on randomly choosing a transition function, permuting its output values, and rearranging the Boolean function according to this permutation. The output given by the program is the number of times that the original attractors are found in the perturbed networks.

**Figure 1.**
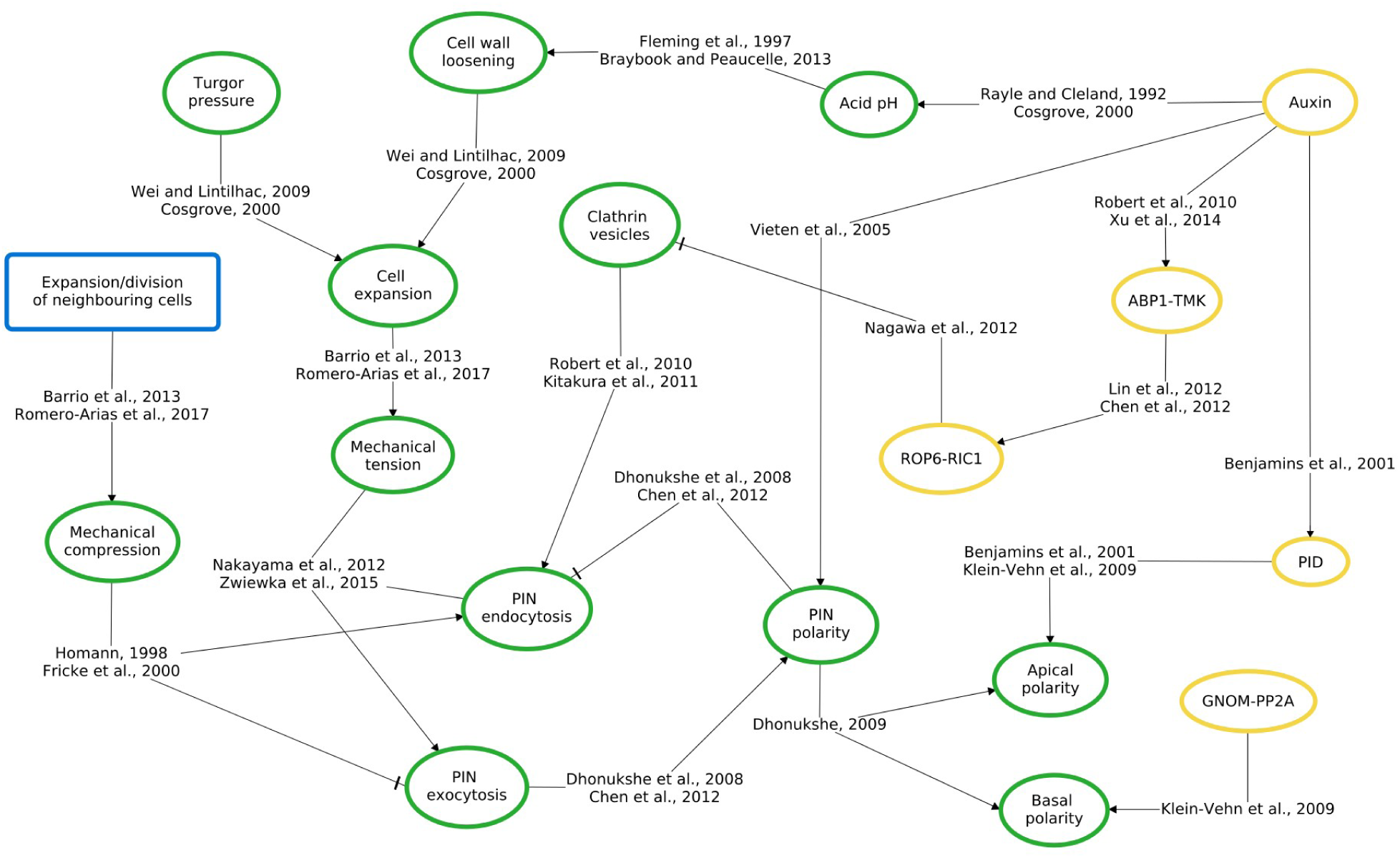
A biophysical dynamic module for the polarisation of PIN auxin efflux transporters. Arrows indicate positive regulation and flat arrows denote negative interactions. Circles denote cell autonomous processes and the rectangles relate to those of tissue scale events. Yellow circles indicate molecular interactions and green ones are cellular scale processes.

**Figure 2.**
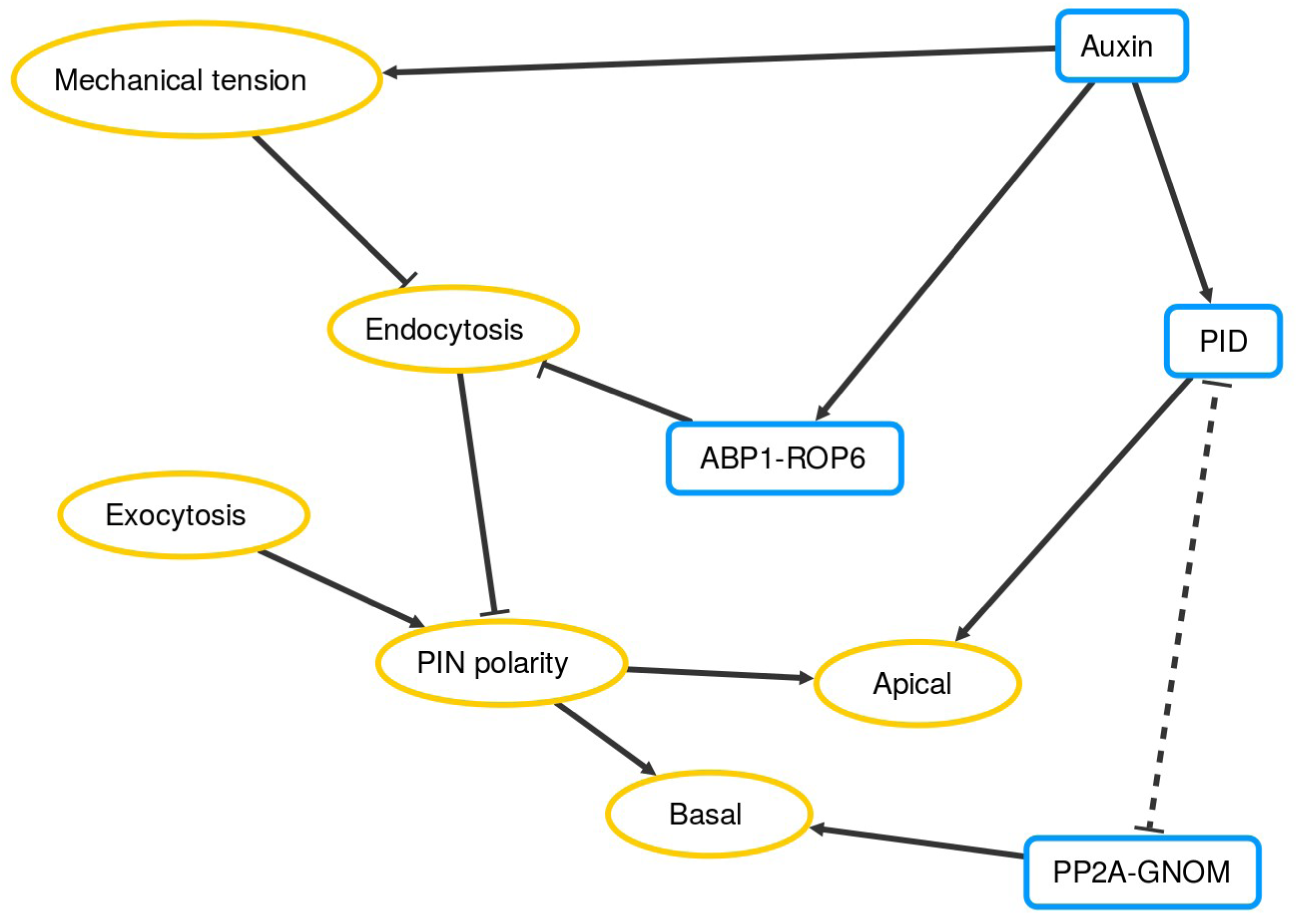
Cell-autonomous network of the biophysical dynamic module for PIN polarisation. Arrows indicate activations and flat arrows denote inhibitory interactions. Yellow circles indicate cellular-scale processes and blue rectangles represent molecules. Dotted lines are for interactions that are not grounded on experimental evidence but we postulated in this study.

### Continuous model

A more versatile and realistic approach to model the inner mechanisms of PIN polarisation is the use of regulatory networks involving state variables and parameters varying within a continuous range. In this approach, the discrete logical inputs are replaced by differentiable continuous functions 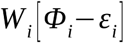 These functions display a step-like (logistic) behaviour which depends on a continuous realisation 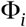 of the discrete proposition 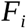 and a threshold level 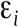 (usually, ε_*i*_ = 0.5). When 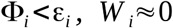 while for 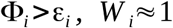 (Mendoza and Xenarios, 2006; Villarreal et al. 2012). In order to translate the discrete Boolean functions into continuous functions we used continuous logical rules to develop a Boolean algebra in the following way (Azpeitia et al., 2014):

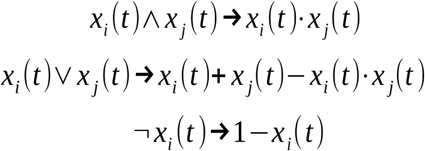

These rules satisfy Boolean axiomatics, and are equivalent to those of Fuzzy logics formerly proposed by Zadeh to investigate properties of control systems (Zadeh, 1965). The state variables considered in this approach should be interpreted as relative values with respect to some fiduciary level as before, and they should be considered as a qualitative representation of the system. So, for example, the logic input for the endocytosis node:

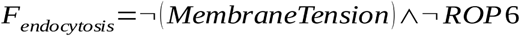

was transformed under the former rules to:

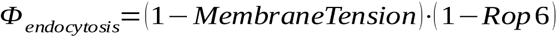

Our approach to study the dynamics of the continuous Boolean network considered that the system is described by a set of differential equations (see Azpeitia et al., 2014):

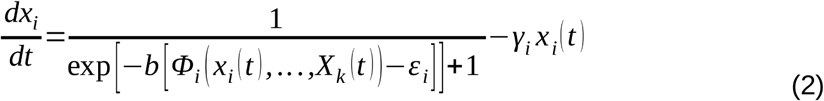

where the logistic input function involves a saturation rate 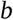, and 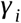 is the decay rate of node *i* activity. For 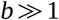 (in this case 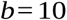) the input function displays a dichotomous step-like behaviour in which the maximum and minimum activation levels of 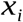 reach values 1 and 0, respectively, permitting a direct comparison with the discrete Boolean model. The set of all Boolean functions and their continuous version can be consulted in Table 1. The simulations of the continuous model were performed using Wolfram Mathematica (version 7.0.0). With the purpose of studying the modifications to PIN polarisation dynamics arising from the continuous description, the numerical solutions of the set of differential equations were obtained by introducing the activation-inhibition patterns of the WT cyclic attractors of the discrete model as initial values. We varied the decay rates 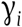 below and beyond of the central value 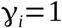 (implicitly considered in Boolean approximation). This variation represents a loss- 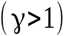 (LF) and gain-of-function 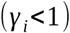 (GF) that can be related to mutations and microenvironment changes, or alterations of intrinsic time expression of a given node. These parameter changes allowed to explore the relative contribution of the different factors that regulate PIN polarisation dynamics.

**Table 1.** Set of Boolean functions, their continuous version and the references that support them

| Boolean function | Continuous function | References |
| --- | --- | --- |
| Auxin: auxin | Auxin(t+1) = auxin(t) | *Smith et al., 2006 |
| ABP1: $\neg$auxin | ABP1(t+1) = 1 – auxin(t) | Robert et al., 2010; Xu et al., 2014 |
| ROP6: auxin $\wedge$ $\neg$ABP1 | ROP6(t+1) = auxin(t) [1 – ABP1(t)] | Kitakura et al., 2011; Chen et al., 2012; Lin et al., 2012 |
| MechTen: auxin $\wedge$ $\neg$exo | MechTen(t+1) = auxin(t) [1 – exo(t)] | Rayle and Cleland, 1992; |
|  |  | Fleming et al., 1997; Morris and Homann, 2001; Cosgrove, 2000 |
| Exo: MechTen | $\text{Exo}(t+1) = \text{MechTen}(t)$ | Homann, 1998; Nakayama et al., 2012; Zwiewka et al., 2015 |
| Endo: $\neg \text{MechTen} \wedge \neg \text{ROP6}$ | $\text{Endo}(t+1) = [1 - \text{MechTen}(t)] [1 - \text{ROP6}(t)]$ | Homann, 1998; Nagawa et al., 2012; Nakayama et al., 2012; Zwiewka et al., 2015 |
| PID: $\text{auxin} \wedge \neg \text{PP2A}$ | $\text{PID}(t+1) = \text{auxin}(t) [1 - \text{PP2A}(t)]$ | Benjamins et al., 2001; Klein-Vehn et al. 2008, 2009 |
| PP2A-GNOM: $\text{PP2A-GNOM} \wedge \neg \text{PID}$ | $\text{PP2A-GNOM}(t+1) = \text{PP2A}(t) [1 - \text{PID}(t)]$ | Klein-Vehn et al. 2008, 2009 |
| PINpol: $\text{exo} \wedge \text{auxin}$ | $\text{PINpol}(t+1) = \text{exo}(t) [\text{auxin}(t)]$ | Vieten et al., 2005; Nakayama et al., 2012 |
| ApicalP: $\text{PINpol} \wedge \neg \text{endo} \wedge \text{PID}$ | $\text{ApicalP}(t+1) = \text{PINpol}(t) [1 - \text{endo}(t)] \text{PID}(t)$ | Klein-Vehn et al. 2008, 2009 |
| BasalP: $\text{PINpol} \wedge \neg \text{endo} \wedge \text{PP2A-GNOM}$ | $\text{BasalP}(t+1) = \text{PINpol}(t) [1 - \text{endo}(t)] \text{PP2A-GNOM}(t)$ | Klein-Vehn et al. 2008, 2009 |

(\*) In our cell-autonomous system, auxin is an input.
MechTen: mechanical tension of the cell membrane; endo: endocytosis; exo: exocytosis; PINpol: PIN localisation at the plasma membrane; ApicalP: apical polarity; BasalP: basal polarity.

## RESULTS

### The biophysical dynamic module

Figure 1 shows the interaction network constructed from the literature. It should be noted that, although the interactions between nodes appear to be direct, the edges can represent a direct or an indirect interaction mediated by one or more intermediate factors. In many cases we still lack the experimental evidence to discern between these two possibilities. In this section we described the factors that are part of our biophysical module for PIN polarisation and how they interact.

Our current understanding suggests that different PIN proteins seem to localise to the plasma membrane using similar mechanisms. For example, the inhibition of endocytosis of the apically localised PIN2 in epidermal cells of Arabidopsis root requires the activity of the GTPase ROP6 and its downstream effector, the ROP-Interactive CRIB motif-containing protein 1 (RIC1) (Chen et al., 2012; Lin et al., 2013). This same mechanism is used for the internalisation of PIN1 located at the lobes of pavement cells in the epidermis of the leaf but in this case it is inhibited by ROP2 and RIC4 (Xu et al 2010, 2011). Also, the effects on PIN polarisation by changes in mechanical forces has been observed in the SAM and RAM of tomato and Arabidopsis (Nakayama et al., 2012; Zwiewka et al., 2015). Based on this assumption, we collected experimental information regardless of the variant of PIN carriers and the organ types studied and generalised our module for all the PINs that localise at the plasma membrane. Our aim is to study the regulatory logic that lies behind PIN polarisation.

As plasma membrane-embedded proteins, PINs undergo constant recycling through endo-, exo-, and transcytosis (Dhonukshe, 2009), and can change their polarisation in response to several types of signals, for example, light (Ding et al., 2011), and mechanical forces (Heisler et al., 2010; Feraru et al., 2011; Nakayama et al., 2012; Braybook and Peaucelle, 2013; Zwiewka et al., 2015). It has been shown that the clathrin-mediated internalisation of PIN carriers is important for the dynamics of PIN polarisation (Klein-Vehn et al., 2011; Kitakura et al., 2011; Nagawa et al., 2012). Several experiments have shown that auxin can inhibit the endocytosis of PINs located at the plasma membrane via ABP1 signalling (Paciorek et al., 2005; Robert et al., 2010; Chen et al., 2012; Lin et al., 2012; Xu et al., 2014). This interaction then upregulates the activity of ROP GTPases that stabilises cortical actin filaments at the membrane that inhibit the formation of clathrin-coated vesicles and, thus, PIN endocytosis (Kitakura et al., 2011; Xu et al. 2010, 2011; Chen et al., 2012; Nagawa et al., 2012; Lin et al. 2012, 2013) (Figure 1).

In addition to the inhibition of endocytosis through the ABP1-ROP-RIC pathway, auxin also seems to regulate the dynamics of PIN polarisation through changes in cellular mechanical forces. According to the acid-growth hypothesis, auxin acidifies the cell wall and this activates cell wall loosening enzymes such as expansins (Rayle and Cleland, 1992; Fleming et al., 1997; Cosgrove 2000, 2005). Loosening of the cell wall then allows cell expansion, which increases the mechanical tension of the plasma membrane. Experiments have shown that when the mechanical tension of the plasma membrane increases as a result of cell expansion, the exocytosis rate becomes higher (Homann, 1998). In the opposite way, when the cell's membrane compresses, endocytosis is higher (Homann, 1998; Fricke et al., 2000; Morris and Homann, 2001). These changes in the rates of endo- and exocytosis are supposed to account for the changes in surface area of the cell – there is retrieval of membrane material when the cell contracts and addition during elongation (Homann, 1998). A hypothesis has been put forward that “a homeostatic relationship exists between the plasma membrane tensions and plasma membrane area, which implies that cells detect and respond to deviations around a membrane tension set point” (Morris and Homann, 2001). In line with this, mechanical modulation using osmotic treatments or by application of external forces, for example, have been shown to affect the polarisation patterns of PIN proteins in *Solanum lycopersium* and *A. thaliana* systems (Nakayama et al., 2012; Zwiewka et al., 2015).

Mechanical forces a cell experiences can be positive – i.e., tension –, or negative – i.e., compression. If we look at a cell *i* within a tissue, a positive force results from the intrinsic cellular elongation that increases the tension in the cell membrane and wall. Cells can also be compressed by the elongation and/or division of neighbouring cells (Barrio et al., 2013; Romero-Arias et al., 2017). Our current understanding suggests that, regardless of where they are produced – at the tissue, cell wall or membrane levels – changes in the mechanical forces are sensed by the membrane and they affect PIN polarisation patterns (Heisler et al., 2010; Feraru et al., 2011; Nakayama et al., 2012; Braybook and Peaucelle, 2013; Zwiewka et al., 2015).

It must be stated that, given the qualitative nature of our approaches and the fact that we did not explicitly model a spatial dimension, we left out of the scope of the models presented here the forces that are produced at the tissue level. Because our models intend to explore the cell autonomous processes, the mechanical stress node refers to the intrinsic tensional force of cell elongation. Another important assumption of our models is that PINs are first located to the plasma membrane after their synthesis and they later attain polarisation patterns, as has been suggested by other authors (Dhonukshe, 2009; Klein-Vehn et al., 2011). Based on this, we defined a node for membrane-located PINs after their synthesis, named PIN polarisation, and two other nodes, the basal and apical polarisations, for the differential distribution patterns (Figure 1 and 2).

After their localisation to the plasma membrane, the phosphorylation state of PIN carriers determines the polarisation patterns at the apical or at the basal sides of the membrane. In the root of Arabidopsis, the PINOID (PID) protein serine/threonine kinase phosphorylates PIN2 located at the membrane, which inhibits recruitment of PIN2 to the basal membrane of the cell (Klein-Vehn et al. 2008, 2009) (Figure 1). On the other hand, the serine/threonine protein phosphatase (PP2A) dephosphorylates PIN2, enabling the protein to be recruited by the GDP/GTP exchange factor for small G-proteins (GNOM) towards the basal membrane (Michniewicz et al., 2007; Klein-Vehn et al. 2008, 2009). The activity of these two nodes in our network is what determines the state of the apical and basal polarity nodes. This same antagonistic mechanism between PID and PP2A-GNOM has been described in pavement cells in the shoot, where the knockout (KO) mutation of *PP2A* and the overexpression (OE) of *PID* both caused a relocalisation of PIN1 from lobe to indentations regions (Michniewicz et al., 2007; Li et al., 2011). For simplicity, we comprised the factors PP2A and GNOM in a single node, named the PP2A-GNOM node, as they have been shown to act in concerted manner (Klein-Vehn et al. 2008, 2009).

### Boolean model

The interaction network we modelled is presented in Figure 2 which, as stated before, only considers cell-autonomous processes and some postulated novel interactions that we describe in this section. For the sake of simplicity, this network collapsed related factors and processes into a single node, representing a branch or unit of a pathway. For example, ABP1's influence on the formation of clathrin-coated vesicles via ROP6-RIC1 is a linear pathway that was collapsed into the node ABP1-ROP6. These modifications did not affect the outcomes of the simulations (data not shown).

Although the Boolean functions generated from the experimental evidence recovered general features of PIN polarisation dynamics, they were not sufficient to reproduce every observed behaviour experimentally observed and reported (Table 1 of Supplementary Material 1). Specifically, the antagonistic effects between the GNOM-PP2A node and the PID kinase in determining the basal and apical polarisation patterns were not recovered. A shift of PIN2 polarity from the basal to the apical membranes of PIN2 in cortex cells of the Arabidopsis roots occurs in *GNOM* loss-of-function and *PID* gain-of-function backgorunds (Geldner et al., 2003; Michniewicz et al., 2007; Klein-Vehn et al., 2009). For the polarisation of PIN2 in epidermal cells of the root, the *PID* OE mutant was expected to result in a value of the basal polarity to be zero, however, this was not the case (attractors not shown). As a result, we considered three possible regulatory scenarios – a unidirectional inhibition, either from PP2A-GNOM towards PID or viceversa; and a double inhibition – between the nodes PPP2A-GNOM and PID that are congruent with the available evidence. We systematically tested the three scenarios to elucidate which one could recover the reported antagonistic roles and summarised the results in the Table 1 of the Supplementary Material 1. We found that the double inhibitory interaction between the two nodes reproduced the expected behaviour of PIN polarisation shifts between the apical and basal membranes when we simulated mutations of the *GNOM-PP2A* and *PID* nodes. This hypothesised interaction was included in the final version of our Boolean network as a postulation (dotted lines depicted in Figure 2).

Synchronous updating of the logical rules of all the nodes of the network yielded four attractors for the WT phenotype: two fixed-point and two cyclic attractors of period four (Figure 3). The fixed-point attractors correspond to the conditions in the system in which there is no auxin, no PIN transporters and thus, no PIN polarity. In contrast, in the cyclic attractors (Figure 3), the polarity of PIN transporters oscillates and so do the apical and basal localisations. The two cyclic and fixed-point attractors differ from each other in that they represent the reciprocal state – i.e., apical or basal polarisation.

**Figure 3.**
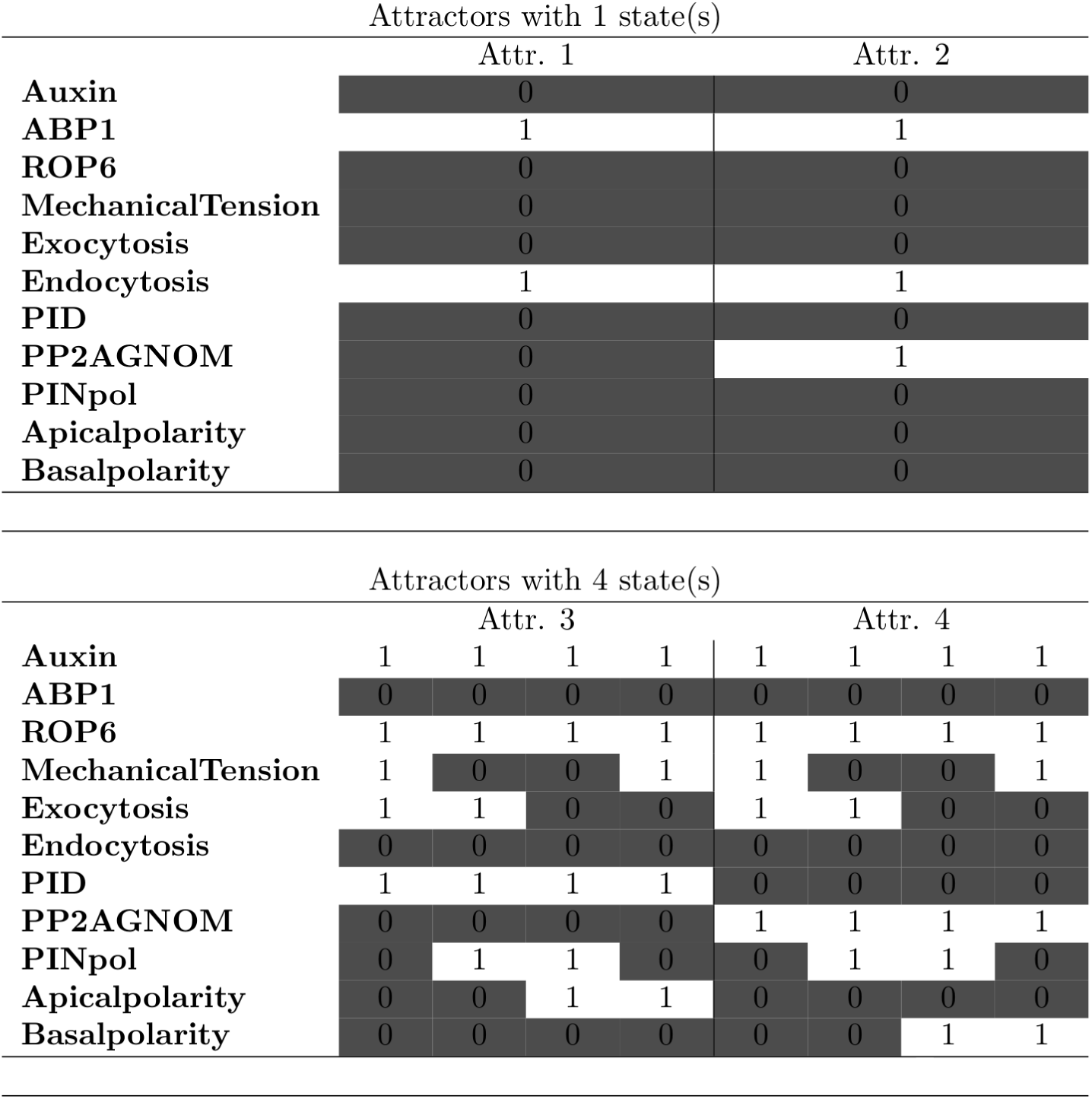
Boolean attractors for the WT condition. The two fixed-point attractors correspond to a condition where the system is depleted of auxin. In contrast, the cyclic attractors present the auxin node active and they have a period of four showing oscillations in the activation state of the mechanical force, exocytosis, PIN polarity and apical/basal polarity nodes.

The direct regulators of the PIN polarity node are the exocytosis, endocytosis and auxin (Figure 1 and 2). Of these three, the activation state of the endocytosis and auxin remains at the same value through the cyclic attractor, 1 and 0, respectively, while the state of the exocytosis oscillates. Note that when a pool of PIN proteins is provided, for example by auxin-dependent gene expression (Vieten et al., 2005), the PIN polarity node is activated one time step after the exocytosis node. This suggests that exocytosis plays a major role in the dynamics of PIN polarisation compared with the other three direct regulators of the PIN polarity node (Figure 3).

Robustness of the model was tested by constructing 1000 copies of the network and perturbing the functions by means of the “shuffle” method that has been described in the *Boolean model* subsection of the Methods. We performed this simulation ten times and summarised the results in Table 3 of the Supplementary Material 1, which shows that in all our simulations the four WT attractors were always recovered.

As a second test to our network construction, we performed simulations of KO and OE mutants for each node of the network and compared the activation configuration reached by these attractors with the reported experimental behaviours. In Table S1.2 of the Supplementary Material 1, we presented a comparison between the published experimental behaviours and the activation state of the nodes of the network reached in the attractors for the WT, KO, and OE simulations. We interpreted that the expected behaviour was recovered when the attractors reached by the mutant was changed from the WT attractors and the change was in agreement with the cited experimental data. As it can be observed in the Table S1.2, most of the attractors of our Boolean model qualitatively recovered expected behaviours.

In Figure 4 we show a comparison of the attractors reached by the KO and OE simulations for the *ROP6* and the increase/decrease of the mechanical force of the plasma membrane. In the case of the OE and KO *ROP6* mutants (the panels A and B of figure 4), the system also attained two fixed point attractors and two cyclic attractors as in the WT. The OE attractors of *ROP6* presented no differences with respect to the WT attractors (see Discussion below). On the other hand, the KO of ABP1 and OE simulations of *ROP6* both behaved in a reciprocal way, which was expected according to what has been experimentally reported (Chen et al., 2012) (Figure S2.4 and S2.7, respectively, of Supplementary Material 2).

**Figure 4.**
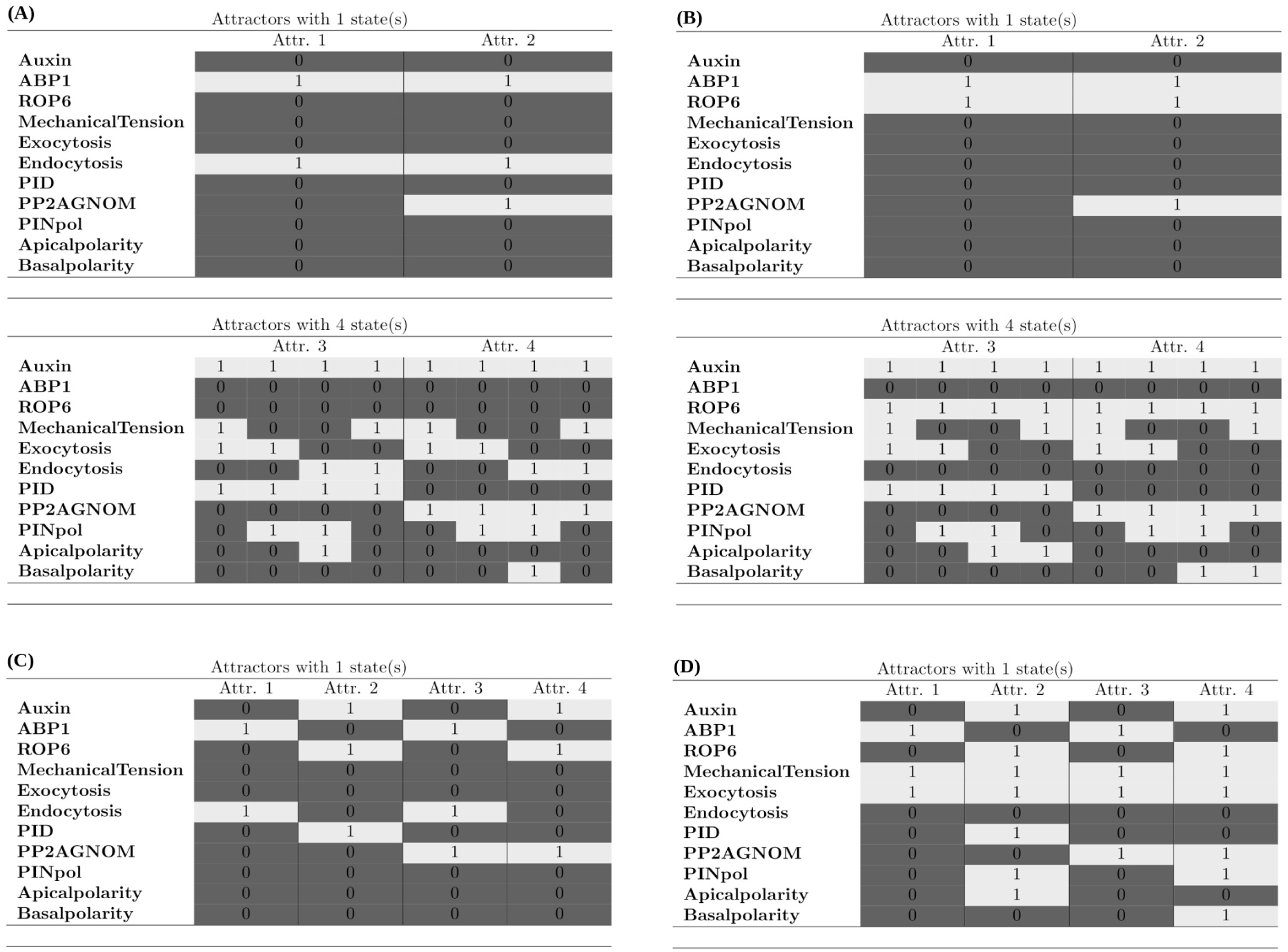
Boolean attractors for mutant conditions. (A) *ROP6* loss-of-function; (B) *ROP6* gain-of-function; complete loss of membrane mechanical force (C); and (D) constant membrane stress. In panels A and B the attractors are similar to those reached in the WT phenotype while, contrarily, in panels C and D the system attains four completely different attractors than in the WT, suggesting that the relative effects of the mechanical force in the polarisation of PIN dynamics is more important than that of the GTPase ROP6.

Although the attractors of the WT and *ROP6* KO simulations are similar, in the case of the cyclic KO attractors the number of time steps in which basal and apical polarities are active is reduced to a single one (Figure 4A). It has been reported that the loss-of-function *ROP6* mutants show an increase in PIN2 internalisation in *A. thaliana* roots (Chen et al., 2012). Because of the discrete nature of the Boolean model, we could not make a direct comparison with the results reported by Chen and coworkers (2012) of a decrease/increase internalisation of plasma membrane located PINs. However, we considered that the reduction in the number of time steps that the apical/basal polarisation of PIN transporters remains active for *ROP6* mutants in our simulations seemed to be in the same direction of the reported behaviour.

For the increased and decreased simulations of the mechanical stress, the system acquired four different fixed-point attractors in contrast with the WT (Figure 4C and D). The most relevant results are those associated to the complete loss of mechanical forces of the plasma membrane (Figure 4C). It can be observed that in two of the four attractors neither the polarity nor the apical/basal nodes are activated, not even when the auxin node is active. This is a more drastic effect on the dynamics of PIN polarisation than the one observed with the *ROP6* KO attractors, suggesting that the role of the mechanical force of the membrane on PIN polarisation dynamics is more important than that of the ROP6-GTPase. This same effect is observed in the decrease and increase simulations of exocytosis (Figures S2.10 and S2.11, respectively, of Supplementary Material 2). On the other hand, this is not the case for endocytosis as the attractors reached in the Boolean simulations of increased/constant endocytosis of PINs (Figure S2.13 of Supplementary Material 2) show that an increase of PIN endocytosis has a stronger effect on its apical/basal distribution than in their localisation at the plasma membrane. A similar observation holds for the contribution of *ABP1* OE (Figure S2.5 of Supplementary Material 2).

### Continuous model

The dynamic analysis in the continuous approximation was performed by assuming that the initial state values were those found for the attractors of the discrete approximation, allowing a straightforward comparison of the system behaviour in both approaches. Particularly, the initial values of the cyclic attractors were considered: auxin = 1, ABP1 = 0, ROP6 = 1, mechanical tension = 1, exocytosis = 1, endocytosis = 0, PID = 1, PP2A-GNOM = 0, PIN polarity = 0, apical polarisation = 0, and basal polarisation = 0.

We observe in Figure 5 the result of the simulation for the WT phenotype when. In this case the value of the PIN polarity, apical polarisation, the membrane mechanical stress and exocytosis stabilise around a value of 0.5 after a transient oscillatory regime. On the other hand, the GTPase ROP6 and the activity of PID remain stable at one, while all other nodes remain inactive. As in the Boolean case, in this continuous version the apical and basal polarisation attractors are also reciprocal from each other (data not shown). It must be stressed that the values of activity level are given in arbitrary units and only reflect the relative contribution of the factors that were taken into account in this study.

**Figure 5.**
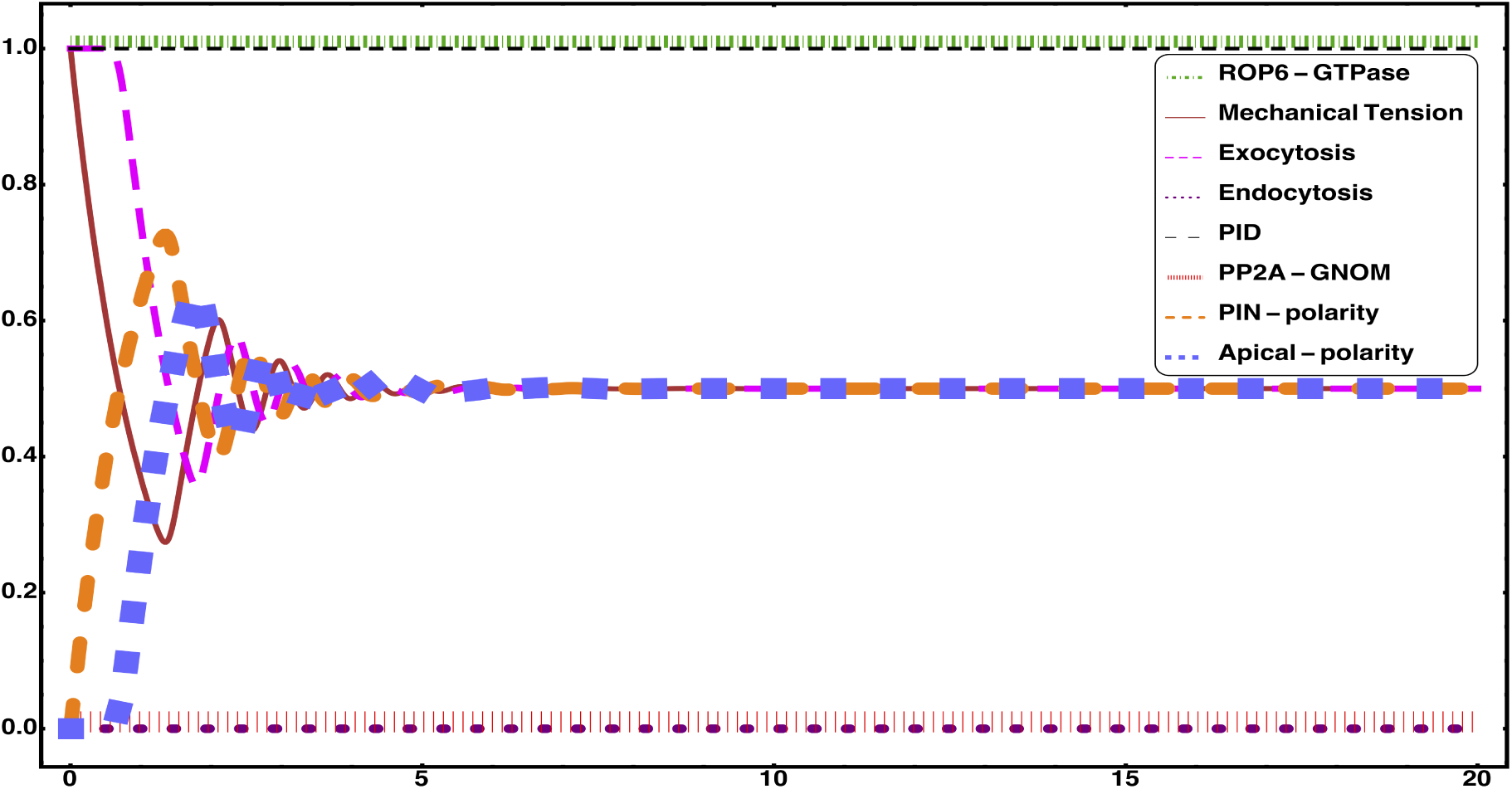
Continuous behaviour of the biophysical dynamic module for PIN polarisation in the wild type. This simulations were performed with a value of the Υ_*i*_, the decay rate of activity, being 1.

As mentioned before, in order to study the modifications of PIN polarisation dynamics that arise from differences on the timing of expression of the network elements, we performed an exploration based on the variation of the decay rate Υ_*i*_ beyond and below the initial value Υ_*i*_== 1. In the loss-of-function simulations we found that, for some nodes, the dynamics of the system was modified with respect to the unaltered behaviour when the corresponding decay rate was augmented by 10 per cent of the initial value. This was the case for the auxin, the membrane mechanical tension, exocytosis, and the PID nodes (Figure 6). On the contrary, the value of γ for other nodes had to be increased up to seven times the initial value in order to obtain a decrease of PIN polarisation. An example of the latter was the simulation for the loss-of-function of *ROP6* (Figure 7A). In this mutant background, we found that when the endocytosis increased rapidly in the first time steps and similarly the apical polarisation which later decreased and stabilised around a value of 0.2 (Figure 7A). This decreased polarisation is in line with what has been experimentally reported (Chen et al., 2012; Lin et al., 2012).

**Figure 6.**
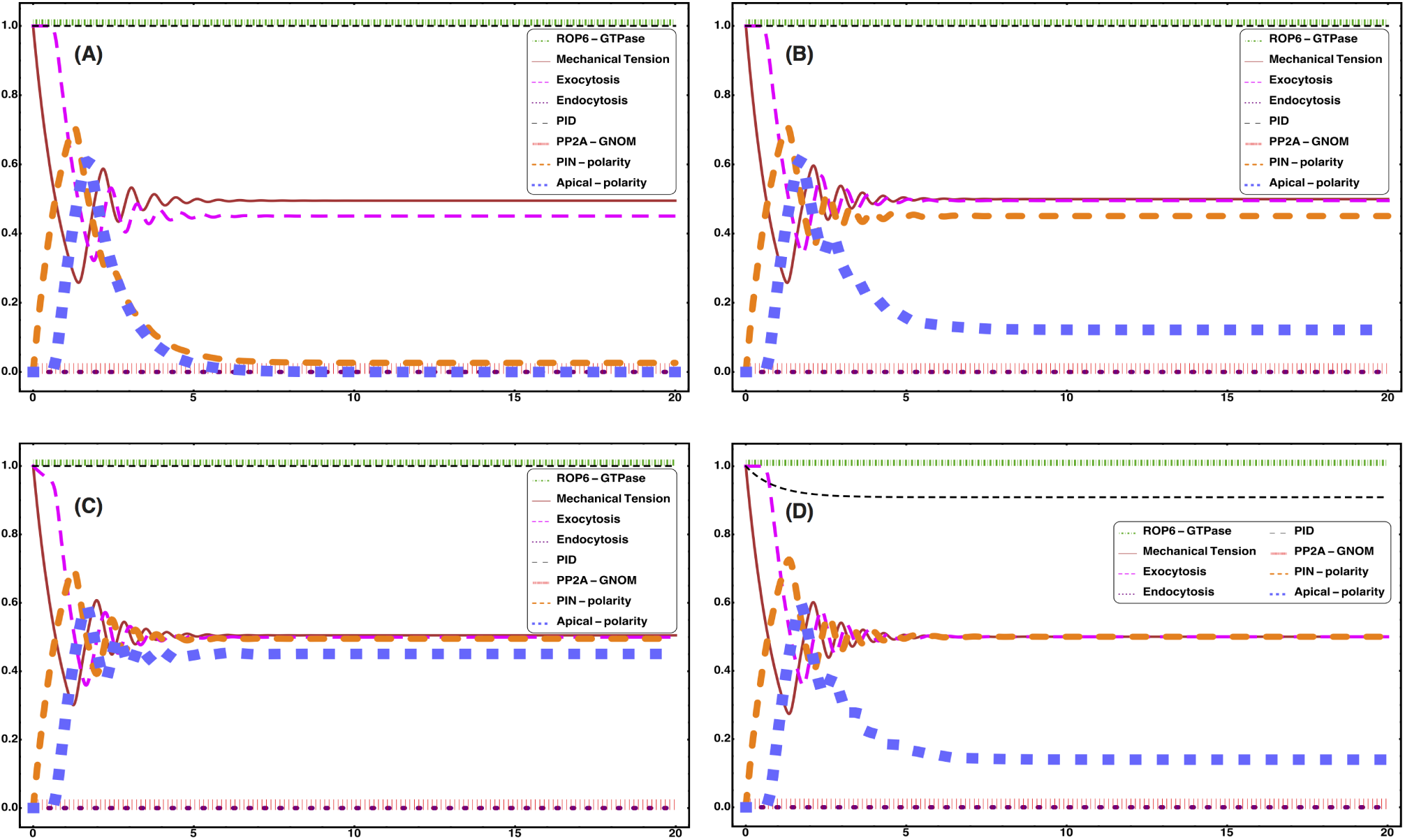
Increases by 10 percent in the value of Υfor some factors of the biophysical module can modify the polarisation dynamics of PIN transporters. We here show the simulations when Υ=1.1 for (A) auxin; (B) the mechanical tension; (C) exocytosis; and (D) PID.

**Figure 7.**
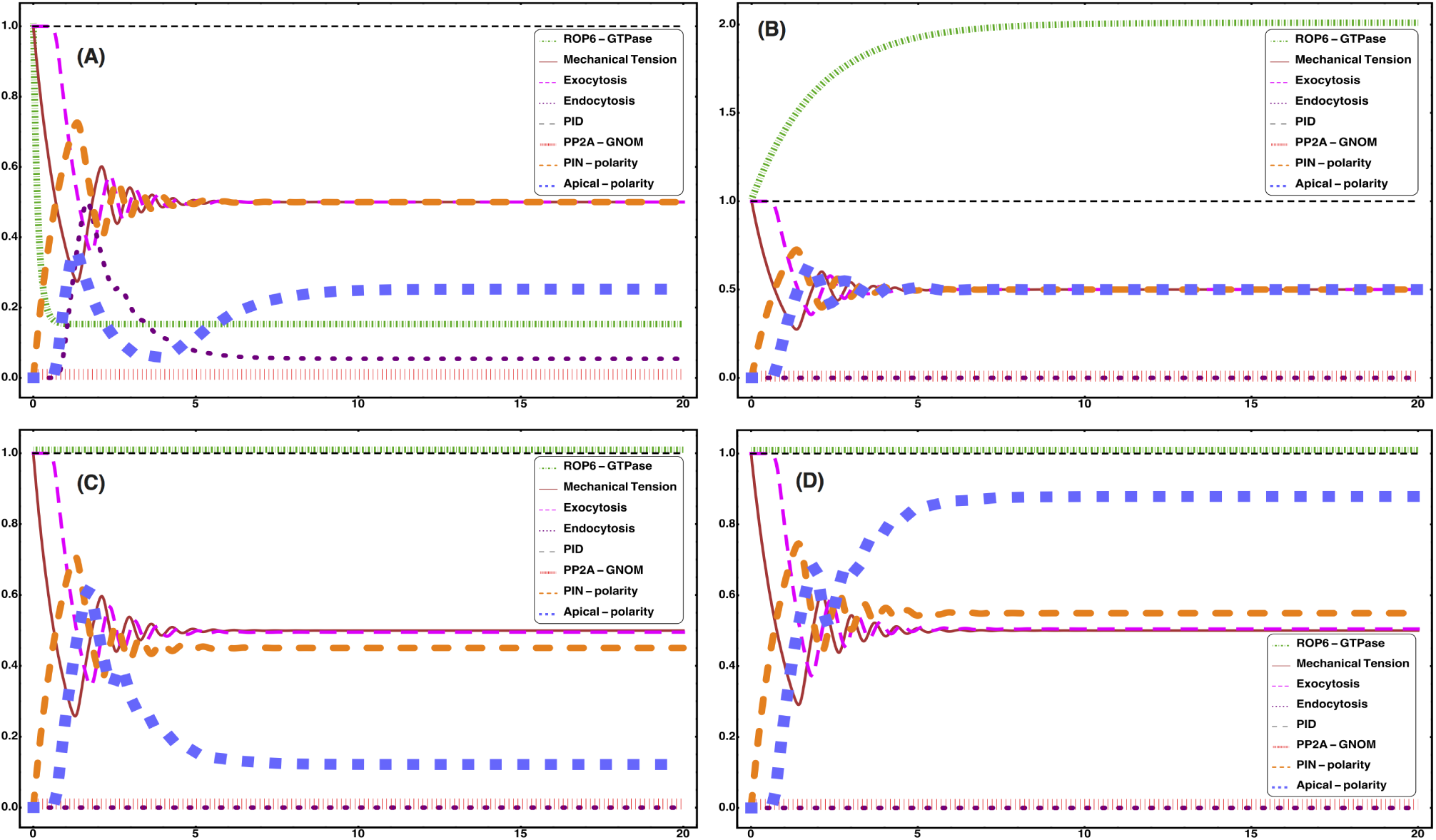
Comparison of the effects of mutations of *ROP6* and increase/decrease of the plasma membrane mechanical tension. Panels A and B account for the *ROP6* loss-of-function and gain-of-function simulation mutations, respectively. In C the membrane mechanical tension has been decreased while in D it has been increased.

In the case of the gain-of-function or increase of activity simulations, a decrease of 10 per cent of the initial decay rate for the mechanical tension was sufficient to see an increase in the value at which the PIN polarity stabilised (Figure 7D), but this was not the case for the ROP6 GTPase. For ROP6, although a wide range of decay rate values were explored (0.1 – 0.9) the simulations never recovered experimentally observed results of an increased polarisation of PIN transporters at the plasma membrane (Chen et al. 2012; Lin et al., 2012) (Figure 7B). As it was shown in the Boolean simulations, the alterations of the γ parameter for the mechanical stress always result in more drastic changes, with respect to the WT phenotype, than those of *ROP6* (Figure 7). In the Supplementary Material 3 we present a complete list of the gain-of-function and loss-of-function simulations of the continuous model for all the nodes of our biophysical module, with 0.5 and 1.5 for γ_*i*_, respectively.

The results depicted in Figure 7 led us to hypothesise that mechanical tension of the plasma membrane is comparatively more influential for the dynamics of PIN polarisation than ROP6 GTPase. We tested this hypothesis first by simulating a condition in which the membrane has a high stress and the activity of the ROP6 is lowered. A second test was simulated with the reciprocal situation – a low mechanical stress in the *ROP6* gain-of-function mutant. In the first case we observed that the effect of the lower *ROP6* activity on PIN polarisation is lost and, even reverted, with an increase of the membrane stress (Figure 8A). In the opposite simulation the effects of a lower mechanical tension on PIN polarisation were not altered by a higher activity of ROP6 (Figure 8B). These results support the hypothesis that mechanical factors can override the effect of changes on key molecular factors involved in PIN polarisation (Zwiewka et al., 2015).

**Figure 8.**
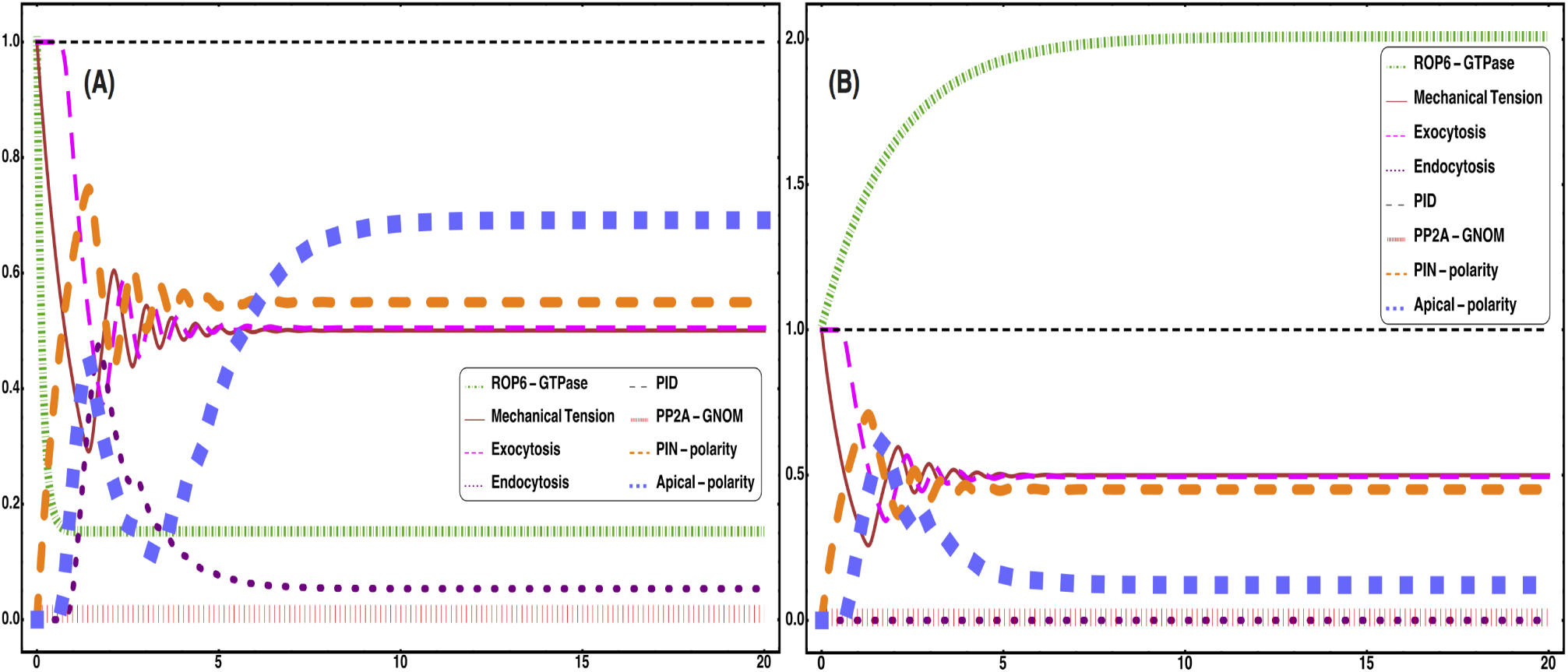
Mechanical stress of the membrane can override the effects of *ROP6* mutations. (A) Mechanical tension of the membrane is increased in a *ROP6* loss-of-function background and (B) decreased mechanical tension in *ROP6* gain-of-function backgorund.

Interestingly, unlike the Boolean attractors, in this continuous model the expected behaviours of *ABP1* gain-of-function and the increased rate of endocytosis were not recovered (Table 2 of Supplementary Material 1, and Figures S3.3-3.4 and S3.11-12 of Supplementary Material 3). However, the behaviour of the dynamical system for the *ABP1* loss-of-function mutant and *ROP6* gain-of-function are reciprocal, just like for the Boolean attractors. As is was expected according to the experimental data, the perturbations of the decay parameter for auxin were different from the original behaviour (Benjamins et al., 2001; Vieten et al., 2005; Smith et al., 2006).

## DISCUSSION

Dynamical models have proven to be a useful tool to address the problem of how plants generate gradients of auxin (Newell et al., 2008; Jönsson et al., 2012). However, these models have mostly focused on the role of molecular mechanisms in the dynamics of PIN polarisation (for a review of several of these models see Berkel et al., 2013). As studies of the biomechanics of plant development have uncovered (Uyttewaal et al., 2012; Barrio et al., 2013; Hernandez-Hernandez et al., 2014; Landrein et al., 2015; Romero-Arias et al., 2017), it has become evident that non-molecular factors must be integrated in these models to understand the generation of auxin gradients and, in fact, whole morphogenetic patterns (Newell et al., 2008; Heisler et al., 2010; Nakayama et al., 2012). The dynamical patterning module concept developed by Newman and coworkers (2009) allows to dissect how morphogenetic patterns emerge from the interactions among molecular, cellular, and biomechanical factors. We used this theoretical framework to propose a biophysical dynamic module for PIN polarisation that integrates factors of different nature, and their interactions. We modelled this biophysical module as either a discrete or continuous network to explore the general regulatory logic behind PIN polarisation dynamics in a single cell. The dynamical patterning framework assumes that the role of physical forces is more or less autonomous from the effects of genes and molecules. The results of both our models are in good agreement with this assumption.

It is considered that after their synthesis, PIN transporters are mobilised to the plasma membrane in a uniform way and, later, differential rates of endocytosis on the different sides of the membrane render an anisotropic distribution (Dhonukshe, 2009; Klein-Vehn et al., 2011). This could be thought of as a two-step mechanism of PIN polarisation. Our simulations of both the continuous and discrete models point to a differentiation between the factors and processes whose contribution is more relevant to establish PIN polarisation patterns from those that locate them at the plasma membrane in a uniform manner. For example, the mechanical stress of the plasma membrane could be considered as a regulator of the uniform localisation of PINs because it affects both the endo- and exocytosis processes. While ABP1, ROPs and RICs could be considered as strong contributors to establish PIN polarisation patterns as they only regulate plasma membrane PIN endocytosis. This implies that studies on PIN polarisation dynamics should look at whether the genetic, chemical or mechanical treatments have an impact on the overall localisation of PINs at the plasma membrane, their polarisation patterns or both.

Although we still lack detailed knowledge about the factors that initially localise PINs at the membrane, an example of the factors involved in the second step is the GTPase ROP6 which is responsible for inhibiting the endocytosis of PIN2 located at the membrane of epidermal cells of the root in Arabidopsis (Chen et al., 2012; Lin et al., 2012). ROP6 and other members of the family of GTPases - such as ROP2 in the pavement cells of the leaf epidermis (Lin et al., 2013) - have not been yet related to the initial localisation of the PINs at the plasma membrane. In pavement cells, ROP6 and ROP2 are activated by extracellular auxin and they promote the assembly of cortical actin filaments through the downstream effectors RIC1 and -4, respectively, which inhibit the formation of clathrin-coated vesicles (Chen et al., 2012; Lin et al. 2012, 2013). This mechanism of action requires that ROP GTPases be located at particular domains of the plasma membrane. For example, ROP2 and RIC4 locate at the lobes of pavement cells and ROP6 and RIC1 locate at the indentations (Xu et al., 2010; Lin et al., 2013), while in pollen tubes ROP1 localises at the tips to promote spatially confined cell expansion (Zhou et al., 2015). This specific localisation of ROP action supports a categorisation of ROP-GTPases to the factors that affect the polarisation of PIN transporters rather than its initial localisation at the plasma membrane.

Although we are aware that this categorisation may not be so straightforward in nature, we consider that the distinction can help uncover the relative contribution of different factors to the polarisation dynamics of PIN transporters. For example, the mechanical stress of the plasma membrane can regulate both endo- and exocytosis (Homann, 1998; Morris and Homann, 2001). Several authors have made the observation that tensile stresses can promote exocytosis and inhibit endocytosis and viceversa (i.e., compression inhibits exocytosis and promotes endocytosis) (Homann, 1998; Morris and Homann, 2001; Nakayama et al., 2012; Zwiewka et al., 2015). Because the localisation of PIN transporters is dynamic, their localisation at the plasma membrane responds to the changes in mechanical forces regardless of the scale at which they were induced (i.e., the plasma membrane or cell wall at the intracellular level or the tissue level) (Heisler et al., 2010; Nakayama et al., 2012; Braybook and Peaucelle, 2013; Zwiewka et al., 2015). Among all the different factors and processes considered in this study, the effects of the changes in the mechanical tension were the most drastic in both our models with respect to the effects of other alterations of the components of our biophysical module. This suggested a significant role of the mechanical tension for the dynamics of PIN polarisation.

We have assumed that the mechanical signal to which cells respond by activating/inhibiting endo- and exocytosis is the mechanical tension. However, it has not been experimentally proven whether cells localise their PINs towards the maximal axis of the mechanical tension or towards the maximal strain axis. Model simulations show that morphogenetic feedbacks that are based on the mechanical tension signal can render complex patterns in a more robust way than those generated responding to the mechanical strain (Bozorg et al., 2014). This aspect of mechano transduction needs to be further studied. Another question that is pending to be answered concerns the definition of the molecular components that transduce the mechanical forces and or strains. The receptor-like kinase FERONIA is a transmembrane protein with an extracellular domain and one candidate factor that mediates mechano transduction (Shih et al., 2014). The loss-of-function mutant *fer* shows altered cytosolic Ca^2+^ concentrations during bend experiments and a reduced upregulation of touch-inducible genes (Shih et al., 2014). The T-DNA insertion mutant also reduces the level of activativated ROP-GTPases and generates root hairs that are either shorter than the WT or collapse after emergence (Duan et al., 2010). It has not been yet explored whether the mutation of FERONIA is capable of impairing PIN polarisation patterns and how.

It has been established that PID phophorylates PIN proteins at the plasma membrane (Klein-Vehn et al., 2009), while the ARF-GEF GNOM factor participates in the exocytic sorting of proteins (Geldner et al., 2003). To be recruited to the GNOM-dependent pathway, PINs must be dephosphorylated by the PP2A phosphatase (Klein-Vehn et al. 2008, 2009). The loss-of-function mutation of *PID* enhances basal polar targeting of PINs, while the *PP2a* mutation has the opposite effect (Klein-Vehn et al. 2008, 2009). When we used only this evidence to generate the Boolean functions of our network and simulated the loss- and gain-of-function mutants of *PID* and *PP2A,* respectively, we did not recover the antagonistic effects on polarisation that have been reported experimentally. Subsequently, we postulated a double inhibition between the PP2A-GNOM and PID nodes. We suggest that the explanation of the inhibitory effect between the basal and apical polarisations could be elucidated by studying how the phosphorylated state of PIN proteins is perceived. According to the experimental evidence, the phosphorylation state of PINs is important not only to determine the polarisation but it can also affect their auxin transport activity (Zourelidou et al., 2014).

Both our discrete and continuous models reproduced expected behaviours when we simulated loss- and gain-of-function mutants, except for the case of the gain-of-function of *ROP6*. It has been demonstrated that several ROPs act in an orchestrated manner to create cellular patterns. For example, ROP1 in the pollen tubes regulates tip growth by activating the different downstream effectors, RIC1, 3 and 4 (Zhou et al., 2015). In the pavement cells ROP6 and RIC1 promote the bundling of microtubules at the necks while ROP2 and RIC4 promote actin assembly at the lobes (Lin et al. 2012). Based on the experimental results from the work of Chen and coworkers (2012) one can see that the effect of mutations of *RIC1* is bigger than that of *ROP6.* A possible explanation for why our models could not recover the expected behaviour of the *ROP6* gain-of-function background, is that there might be a second regulator of RIC1, most probably another member of the ROPs family.

Another possible explanation for the mismatch between experimental works and our simulation's results is related to the use of the drug Brefeldin A (BFA) as an indicator of endocytosis. In the studies that have used BFA, the authors concluded that PINs that are present in BFA vesicles come from endocytosis from the plasma membrane (Paciorek et al., 2005; Robert et al., 2010; Chen et al., 2012). However, BFA is known to interfere with the exocytic protein sorting from the endoplasmic reticulum to the Golgi apparatus (Donaldson et al., 1992; Lusching and Vert, 2014). In the recent publication by Jásik and coworkers (2016), the authors have demonstrated that PINs contained within BFA vesicles are newly synthesised proteins, which strongly suggests that PINs contained within BFA vesicles are not a good indicator of PIN endocytosis. BFA can thus be a powerful tool to examine the plasma membrane location of newly translated PIN proteins, rather than the PIN endocytosis.

BFA has also been used in several studies that conclude that the putative membrane-localised auxin receptor ABP1 is necessary for the auxin-mediated inhibition of PIN endocytosis. These studies observed an increase in the PIN2-GFP signal at the plasma membrane as a consequence of this treatment in the *ABP1* gain-of-function background (Paciorek et al., 2005; Robert et al., 2010; Chen et al., 2012). Neither of our models recovered such an increase of PIN polarisation nor apical/basal distribution in *ABP1* loss-of-function or *ROP6* gain-of-functio*n*. The study by Jásik and coworkers (2016) also demonstrated that chemical analogues of auxin that supposedly inhibit PIN endocytosis through the ABP1 pathway did increase the PIN density at the plasma membrane. Furthermore, null mutants of *ABP1* recently constructed with the CRISPR technique did not result in such an effect (Gao et al., 2015). The results of our simulations are in line with the published experimental evidence (Gao et al., 2015; Michalko et al., 2015; Jásik et al., 2016), which calls for a reflection on the genetic and chemical methods used to uncover auxin action in PIN polarisation dynamics, and to elucidate whether the auxin-ABP1-ROP6 signalling pathway is mediating PIN endocytosis or not.

A novel prediction of our simulations is the role of the mechanical stress of the plasma membrane which can surpass the effect of the ROP6 GTPase in the process of PIN polarisation. As we have previously mentioned, the mechanical stress is the only factor that regulates both the endo- and exocytosis processes. In all the simulations of loss- and gain-of-function of the factors included in our biophysical module the mechanical tension always produced more drastic changes when compared with the effects of other factor's changes. In fact, in the continuous model we had to increase and decrease the value of the γ parameter several more units for the ROP6 than for the mechanical tension in order to obtain comparable results. Recently, Zwiewka and coworkers (2015) found that the effects of osmotic stress, that are supposed to alter the mechanical stress of the membrane, predominate upon the effects of molecular factors such as auxin. When the osmotic potential of the membrane increases, water moves into the cell, increasing the turgor pressure and cell size. This, in turn, increases the mechanical tension of the plasma membrane and, as a consequence, the cell activates the exocytosis of membrane proteins and decreases endocytosis so that an equilibrium at the membrane surface is reestablished (Homann, 1998; Morris and Homann, 2001). The opposite happens when the cell looses turgor pressure and shrinks, activating endocytosis to remove excess membrane and inhibiting exocytosis and further deposition of membrane material (Homann, 1998; Morris and Homann, 2001).

Finally, to test our hypothesis, we performed a simulation of increased mechanical stress in the loss-of-function *ROP6* background and viceversa (a decrease mechanical stress in gain-of-function *ROP6*). In both cases we observed that the effects of the mechanical stress on the rates of endo- and exocytosis as well as in the polarisation patterns of PINs, are predominant over those of the ROP6. These in silico predictions can be experimentally validated in vivo by treating *ROP6* and *RIC1* gain-of-function mutants with hyperosmotic solutions and observe if the increased internalisation (cytoplasmic/plasma membrane ratio) that has been reported for these backgrounds is reversed by the reduction in the membrane tension that results from lower turgor pressure. Another important question to be answered is wether these experiments affect PIN polarisation, the anisotropic distribution patterns in the plasma membrane or both.

## CONCLUSIONS

Our biophysical dynamic module is built upon the up-to-date experimental results on the influencing factors and cellular processes and their non-linear interactions that regulate the polarisation of PIN auxin efflux transporters at the cell level. The modelling and simulations of this module helped postulate that relative influence of the mechanical tension of the plasma membrane can surpass that of the GTPases ROP. Furthermore, we have been able to put forward some novel predictions that can be tested experimentally. Our model's prediction states that the effects of the mechanical stress on PIN polarisation dynamics could revert the decrease and increase in the plasma membrane localisation of ROP6 loss- and gain-of-function mutations.

## ACKNOWLEDGEMENTS

The authors declare no competing interests. VHH and MB acknowledge the graduate program “Posgrado en Ciencias Biológicas” of Universidad Nacional Autónoma de México and Laboratorio de Nacional de Ciencias de la Sostenibilidad. This work was supported by the scholarship support provided by CONACyT to VHH. RAB is grateful for a sabbatical grant from DGAPA and to Prof. Lee Eiden from NIMH for his hospitality during the development of this work. CV was financially supported by project CONACyT 180380. NN acknowledges the University of Edinburgh Chancellor’s Fellowship and the Royal Society University Research Fellowship. JRRA is grateful to CONACyT and FORDECyT Grant No. 265667 for partial financial support during the development of this work. The authors also thank Carlos Galván-Ampudia for valuable comments to the manuscript.

